# A decline in follicle cell function is a major driver of *Drosophila* ovarian aging

**DOI:** 10.64898/2026.02.05.704044

**Authors:** Emily A. Wolfgram, Todd G. Nystul

**Author notes:** Corresponding Author, 513 Parnassus Ave, San Francisco, CA 94143, USA.

## Abstract

The ovary is one of the first organs to lose functionality with age. We found that aging of the *Drosophila* ovary is characterized by an accumulation of phenotypes in the somatic compartment, including failure of the follicle cells to encapsulate germ-cell cysts, an extended S phase, and increased DNA damage. In aged ovaries, follicle encapsulation defects are associated with the lack of a germ-cell cyst checkpoint in early oogenesis. Single-cell RNA sequencing revealed that, across all cell types in the ovary, cells in the follicle lineage have the highest number of differentially expressed genes. Overexpression of *Atg8a*, a key autophagy machinery gene homologous to mammalian LC3, specifically in follicle cells prevents age-associated decline in the follicle epithelium and loss of reproductive capacity. Collectively, these findings demonstrate that genetic manipulation of a small population of ovarian somatic cells is sufficient to improve both cell-autonomous and non-autonomous features of reproductive aging.

## Introduction

Aging is an inevitable biological process that leads to the decline of cellular functionality [1]. Among the organs affected, the ovary is one of the first to lose function, with females reaching menopause at an average age of 51 [2]. While ovarian morphology varies between species, the mechanisms governing oogenesis are highly conserved. From *Drosophila* to higher-order mammals, oogenesis involves the progression of early germ cells into fully developed eggs [3]. There are broad similarities in organ morphology, with many direct parallels between cell types and stages of oogenesis, as well as several conserved features at the molecular and cellular level. Both *Drosophila* and mammalian ovaries have germ cells that undergo multiple rounds of incomplete division, forming multicellular cysts [4]. In the somatic compartment, the follicle cells in the *Drosophila* ovary functionally parallel the granulosa cells in the mammalian ovary. Both follicle cells and granulosa cells directly contact germ-cell cysts and proliferate during oogenesis to maintain a single continuous layer that forms the surface of the growing follicle [5]. In both fly and mammalian ovaries, these cells help guide the process of oogenesis via direct cell-cell communication with germ cells through gap junctions [6,7] and the production of endocrine signals that coordinate the progression of germ cell development with the actions of other cell types [8,9].

The *Drosophila* ovary is comprised of ∼16 strands of ovarioles, with a germarium at the anterior end of each ovariole. The germarium is where oogenesis begins and is host to both the germline stem cells (GSCs) and follicle stem cells (FSCs). At the anterior tip of the germarium reside the GSCs, which asymmetrically divide to self-renew and produce a germ cell. Each germ cell undergoes four rounds of incomplete mitosis, producing a 16-cell germline cyst that moves posteriorly into the germarium at a position called the Region 2a/2b border. At this position in the germarium, a population of follicle stem cells (FSCs), supported by an escort cell (EC) niche, asymmetrically divide to self-renew and produce pre-follicle cells (pFCs) [10,11]. These pFCs will continue to divide and differentiate into polar cells, stalk cells, and main body (MB) follicle cells. When a germline cyst reaches region 3 of the germarium, it buds off, encapsulated by MB follicle cells, and is connected to other encapsulated cysts via stalk cells. The germline cyst continues to grow until one cell is selected as the egg, the other 15 cells becoming nurse cells that support the developing egg. Follicle cells continue to divide and grow to maintain encapsulation of the developing cyst, prior to full egg development [12]. Given the continuous demand for follicle cells to encapsulate and guide the germline cyst through oogenesis, FSCs and pFCs are among the most proliferative cells in adult *Drosophila* [13]. Any stressors that impair FSC proliferation and differentiation could have severe consequences for fertility. Indeed, genetic manipulations that disrupt FSC activity often result in significant morphological defects and, in some cases, sterility. These findings underscore the crucial role of the follicle lineage in *Drosophila* ovarian aging.

The average lifespan of wildtype *Drosophila* is approximately 70 days [14], yet fertility begins to decrease after the first week of adulthood and drops below 10% of the maximum rate by 50 days [15]. The causes of this decrease in fertility are not fully understood but, as in mammals, it involves a decline in germ cell number and is strongly influenced by nutrition and stress signaling pathways [16,17]. Likewise, emerging evidence from studies of both *Drosophila* and mammals is beginning to elucidate the important role of ovarian somatic cell aging in the loss of fertility [18–20]. There are only 2-4 FSCs per ovariole, and yet each ovariole contains several thousand follicle cells, so FSC daughter cells undergo up to eight transit-amplifying divisions before terminally differentiating to meet this demand [13,21]. This makes the FSC lineage particularly susceptible to replicative stress and may make it more vulnerable to other insults that are known to increase with age including stem cell maintenance, reactive oxygen species production, mitochondrial dynamics, and autophagy regulation [1].

In this study, we conducted a phenotypic analysis of aged ovaries and identified cellular and tissue-scale aberrations in the typically well-organized layer of follicle cells. In addition, we performed single-cell RNA sequencing and found a particularly high number of transcriptional differences in the FSC lineage between young and aged ovarioles. We identified *Atg8a*, an upstream effector in the autophagy pathway, as a key regulator of follicle cell aging and reproductive capacity. These results highlight a new avenue for extending reproductive longevity through manipulation of the somatic lineage rather than the germline, which is ultimately passed on to offspring [22].

## Methods

### Fly husbandry and aging time course

Flies were maintained on standard molasses food at 25ºC. The following stocks from BDSC were used in this study: *w1118(x);;* (3605), ;*109-30-Gal4;* (7032), and ;*UAS*-*Atg8a::mCherry-GFP;* (37749). *;;Lamp1::3x-mCherry* was described in Hegedús et al, 2016 [23]. We used FlyBase to find information on phenotypes/function/stocks/gene expression. For all aging assays, virgin females were collected and paired with males and maintained in standard lab conditions until dissections at weekly timepoints. Age was calculated as weeks post-eclosion.

### Immunofluorescence staining and imaging

*Drosophila* ovaries were dissected in PBS and ovaries fixed in 4% PFA + PBS for 15 minutes. Ovaries were rinsed with PBS before blocking for 1 hour in standard block (PBS + 0.2% Triton X-100 + 0.5% bovine serum albumin. Ovaries were incubated in primary antibodies with block overnight on a nutator at 4ºC in dark conditions. The following day, ovaries were rinsed with block for 1 hour before adding secondary antibodies. Ovaries were incubated in secondary antibodies with block overnight on a nutator at 4ºC. The following day, ovaries were rinsed three times with PBS for 5 minutes each. Once the PBS was removed, DAPI-Fluoromont G Slide Mounting Media (SouthernBioTech) was added, and ovaries were stored at 4ºC in dark conditions until mounted on slides. Ovaries were mounted on glass slides in DAPI-Fluoromont G Slide Mounting Media. Primary antibodies used were: mouse anti-Fas3 7G10 (1:100, DSHB), rat anti-Vasa 46F11 s(1:200, DSHB), rat anti-CadN DN-Ex#8 (1:100, DSHB), rabbit anti-pHH3 (Ser10, 1:500, Millipore Sigma), mouse anti-UNC93-5.2.1 (gH2Av, 1:100, DSHB), and rabbit anti-Vkg (1:1000, Dr. Stephane Noselli), rat anti-RFP 5F8 (1:500, Chromatek), mouse anti-Rab7 (1:100, DSHB). The following secondary antibodies were used at 1:500: goat anti-mouse 488 (Life Technologies, A11001), goat anti-rabbit 488 (Invitrogen, A11034), goat anti-rat 488 (Invitrogen, A11006), goat anti-mouse 555 (Invitrogen, A21422), goat anti-rat 555 (Invitrogen, A21434), goat anti-mouse 647 (Invitrogen, A21236), goat anti-rat 647 (Invitrogen, A21247).

Images were acquired with a Zeiss M2 Axioimager with an Apotome unit using Zen 3.5 software. Initial image processing was performed in Zen using the Apotome Raw Convert feature. All further image processing was done in FIJI, and figures were prepared in Adobe Illustrator.

### Image Analysis

Analyses for N-cadherin prevalence, pHH3, EdU, gH2Av, TUNEL, and Lysotracker were performed by hand-counting. Vkg intensity was performed by outlining the germarium with a segmented line and taking the intensity of Vkg signal after background subtraction.

### EdU staining

For EdU experiments, ovaries were dissected in PBS and incubated in 0.75uL EdU (Click-IT Alexa Fluor 555 Imaging Kit (ThermoFisher Scientific, C10338) with 499.25uL PBS at room temperature for 1 hour. Ovaries were then fixed in 4% PFA + PBS for 10 minutes and then rinsed in PBS prior to blocking for 1 hour in PBS + 0.2% Triton X-100 + 0.5% bovine serum albumin. Primary antibodies were then added, and ovaries were stored overnight at 4ºC on a nutator in dark conditions. The following day, ovaries were rinsed in PBS twice and incubated in a reaction mix (430uL 1x Click-IT EdU Reaction Buffer, 20uL CuSO_4_, 1.2uL Alexa Fluor 555 azide, 50uL 10x Click-IT EdU Buffer Additive in deionized water) for 1 hour at room temperature on a nutator in dark conditions. The ovaries were then rinsed twice with PBS and blocked for 1 hour. Secondary antibodies were added with block and ovaries were stored overnight at 4ºC on a nutator in dark conditions. Ovaries were then labeled with secondary antibodies according to the standard Immunofluorescence Staining protocol described above.

### LysoTracker staining

For LysoTracker experiments, ovaries were prepared according to the immunofluorescence staining protocol described above, except that they were incubated in LysoTracker Red DND-99 (ThermoFisher Scientific) at a 1:100 dilution in PBS for 5 minutes prior to fixation, washed with PBS for 5 minutes, and then fixed in 4% PFA + PBS for 10 minutes rather than the standard 15 minutes. Ovaries were then labeled with primary and secondary antibodies according to the standard Immunofluorescence Staining protocol described above.

### TUNEL staining

For TUNEL experiments, the following protocol was adapted from the Click-iT TUNEL Alexa Fluor Imaging Assays 488 kit (ThermoFisher Scientific, C10617). Ovaries were dissected in PBS and fixed in 4% PFA + PBS for 10 minutes and then rinsed in PBS. Cell membranes were permeabilized in PBS + 0.2% Triton X-100 for 15 minutes. Ovaries were rinsed twice with PBS and then primed with TdT Reaction Buffer for 10 minutes. Ovaries were incubated at 37ºC for 1 hour in a TdT reaction cocktail (47uL TdT Reaction Buffer + 1uL EdUTP, and 2uL TdT enzyme). A reaction supermix was prepared and stored at 4ºC (2630uL 1x Click-iT Plus TUNEL Reaction Buffer, 67uL copper protectant, and 3.7uL 488 picolyl azide). Ovaries were rinsed twice with PBS and incubated at 37ºC for 30 minutes, in a Click-iT reaction cocktail (45uL reaction supermix and 5uL 10x Click-iT Plus TUNEL Reaction Buffer Additive), protected from light. Ovaries were then labeled with primary and secondary antibodies according to the standard Immunofluorescence Staining protocol described above.

### Egg Laying Assay

For egg counts, flies were separated into vials containing two females and one male for each genotype, on standard food at 25ºC. After 24 hours, eggs were hand counted through a dissection microscope and recorded. Total egg counts were divided by 2, so that final data represents the number of eggs per female in a 24-hour period.

### Single-cell RNA sequencing prep

Preparation for single-cell RNA sequencing of *w1118(x);;* ovaries at 1 and 6 weeks old followed the protocol established from Meyer et al, 2024 [24]. However, we did not enrich for the anterior region, as this caused the too much cell loss in the aged condition. The single-cell RNA sequencing was performed at UCSF CoLabs using the 10X Genomics Single Cell 3’ kits (v3.3 and v4). Sequencing was performed on Illumina NovaSeq X at the UCSF Center for Advanced Technology (CAT). CAT is supported by UCSF PBBR, RRP IMIA, and NIH 1S10OD028511-01 grants.

### Bioinformatic analysis

Sequencing reads were aligned to the Drosophila reference genome using the 10X Genomics Cloud CLI, which performs Cell Ranger alignment and initial quality filtering. The standard Seurat pipeline (v5) was applied to each individual dataset to filter out low quality cells based on the number of RNA features, number of RNA counts, and percent mitochondria reads (Supplementary Figure 4A-D). Each dataset was individually normalized and scaled. The two replicates for the 1-week datasets were merged using the IntegrateData function. The two replicates for the 6-week datasets were merged using the IntegrateData function and then anchors from the 1-week merged dataset were applied to the 6-week merged dataset. Principal components and clustering neighbor parameters were determined by the strength of the silhouette scores and mean distance within a cluster, as well as identification of known transcript expression for clusters. One cluster was removed because it had very few cells that all came from one condition and one replicate, suggesting that it was likely a small contaminating group of cells. This workflow resulted in a single Seurat object with 20,818 high-quality cells, grouped into 8 clusters, identified by known transcripts. See GitHub repository (https://github.com/NystulLab/WolfgramAgingOvary.git) for full code for building the Seurat object and each figure panel of the data.

### Statistics and data availability

All analysis for single-cel RNA sequencing (Figure 4, Supplementary Figure 3, Supplementary Tables 1-3) were performed in R. The Seurat object and matrix files are available via GEO. All other statistical analysis and graph generation was performed in Python. Boxplots show the median and interquartile values, and whiskers show the range. Barplots show the mean and error bars show the standard deviation. All quantification is a result of at least three or more independent replicates. A Levene test was used to test for equal variance in cases where a Student’s t-test was used. All raw data files for generating graphs and statistical analysis are available via GitHub.

## Results

### Establishment of follicle cell phenotypes associated with aging in the *Drosophila* ovary

To establish hallmarks of follicle cell aging, we assayed for phenotypes in the follicle epithelium at weekly intervals from one to six weeks post eclosion in ovarioles from *w1118(x)* flies. In healthy ovarioles, follicles bud from the germarium in a single file, are surrounded by an intact, single-layered follicle epithelium, and are separated from one another by a single row of stalk cells (Figure 1A,C). Ovarioles with follicles that deviated from this pattern were scored as having a phenotype. The frequency of ovarioles with these phenotypes progressively increased over the six-week time course (Figure 1D, E). We therefore designated the one-week-old ovary as the “young” condition and six-week-old ovary as the “aged” condition. Qualitative assessment of these follicle cell phenotypes produced four categories (Figure 1F, Supplementary Figure 1A-D). The two most prominent categories were epithelial gapping, where follicle cells, identified by the marker Fasciclin3 (Fas3), are absent from the perimeter of the germarium or developing follicle (Figure 1D, yellow short-dashed line), and epithelial dysregulation, in which the follicle cells were misshapen or failed to form into a single-layered epithelium (Figure 1D, white long-dashed box). We also observed Fas3^+^ cells with a rounded morphology consistent with epithelial cell death (Figure 1D, white long-dashed box), and a small number of instances of impaired stalk development (Figure 1D’’-1D’’’, white arrows).

**Figure 1:**
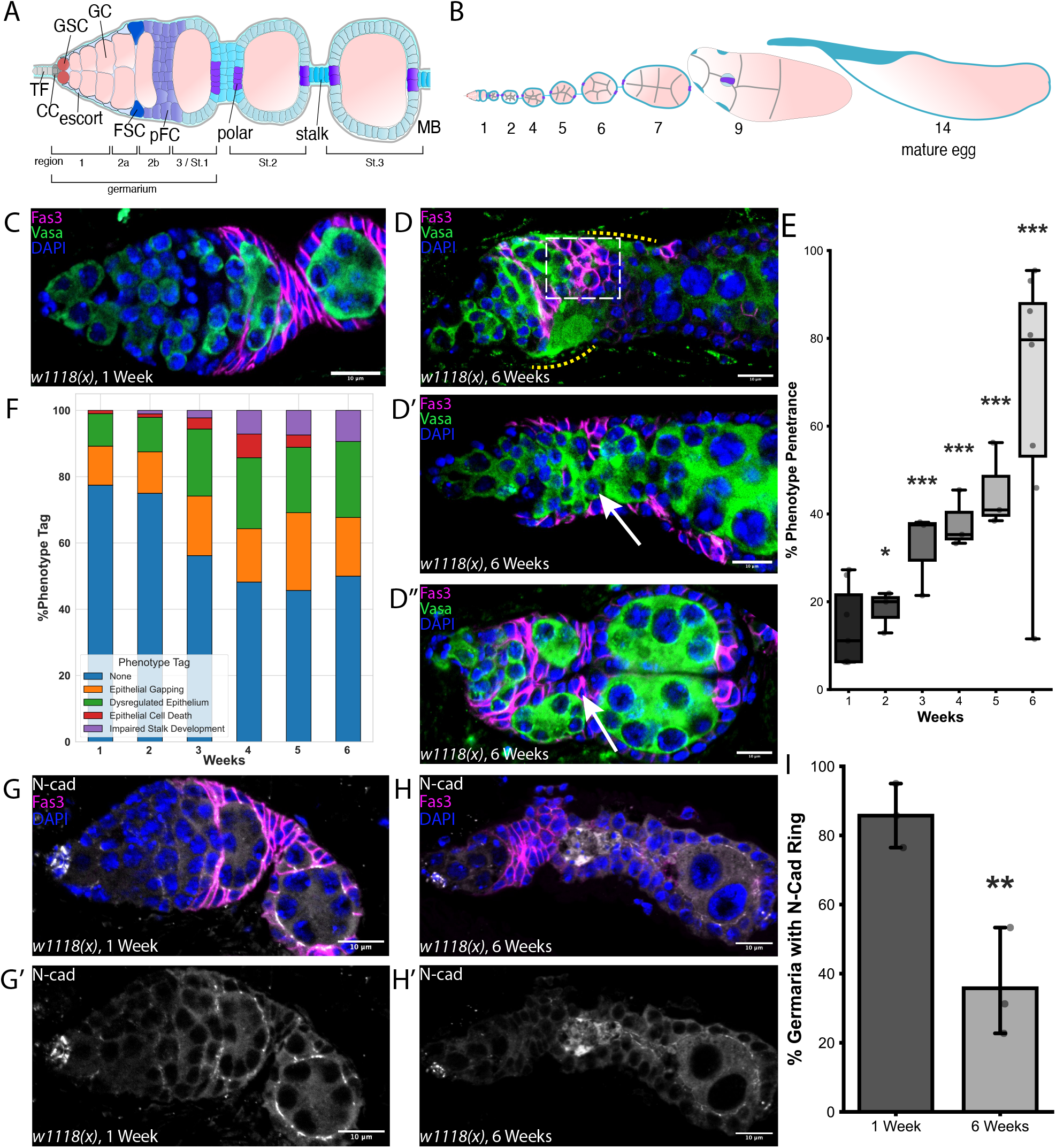
Aging leads to increased follicle cell phenotypes in the *Drosophila* ovary. **(A)** Schematic of the *Drosophila* germarium, positioned at the anterior tip of the ovariole. TF: terminal filament, GSC: germline stem cell, EC: escort cell, FSC: follicle stem cell, pFC: prefollicle cell, MB: main body. **(B)** Schematic of the ovariole. **(C)** Germarium from a *w11118(x);;* fly at 1 week old, stained for Fas3 (magenta), Vasa (green), and DAPI (blue). **(D-D’’)** Representative examples of germaria from w1118(x);; flies at 6 weeks old. Yellow dotted lines show epithelial gapping, white dashed box shows epithelial dysregulation and epithelial cell death, white arrows show impaired stalk development. **(E)** Quantification of the percent of germaria with follicle cell phenotypes at weekly timepoints from 1 to 6 weeks old. n = 298, 88, 73, 43, 64, and 201 germaria, respectively. **(F)** Quantification of follicle cell phenotype tags from 1 to 6 weeks old. n = 98, 88, 73, 43, 64, and 77 germaria, respectively. **(G)** Germarium from a *w11118(x);;* fly at 1 week old, stained for N-cad (grey), Fas3 (magenta), and DAPI (blue). **(G’)** N-cadherin ring forms around developing germ-cell cysts in 1-week-old germaria. **(H)** Germarium from a *w11118(x);;* fly at 6 weeks old, stained for N-cad (grey), Fas3 (magenta), and DAPI (blue). **(H’)** Loss of N-cadherin ring around developing germ-cell cysts in a 6-week-old germarium. **(I)** Quantification of the percent of germaria with the N-cadherin ring around germ-cell cysts at 1 and 6 weeks old. n = 58 and 53 germaria, respectively. Scale bars are 10 µm. ns = not significant, *p<0.05, **p<0.01, ***p<0.001, using Dunnett’s Test (E) and Student’s t-test (I).

The loss of follicle cell integrity in aged germaria prompted us to investigate whether another follicle cell adhesion protein, N-cadherin (N-cad) is also diminished with age. N-cad is highly expressed in prefollicle cells and main body follicle cells from Stage 1 to approximately Stage 5 [10]. N-cad mediates cell adhesion through homotypic interactions at the apical-lateral surface of the cells [25], creating a characteristic “ring” of N-cad signal between follicle cells and germ cells (Figure 1G, G’). We found that this N-cad ring is frequently lost in aged germaria (Figure 1H, H’, I). In contrast, we found no significant difference between young and aged germaria in the level of Collagen IV (*Vkg*) in the basement membrane (Supplementary Figure 2A-C, B’-C’). Likewise, we performed a TUNEL assay to identify apoptotic cells and found that the frequency does not significantly change in aged conditions (Supplementary Figure 2C-E, C’-D’, yellow arrows) [26]. However, the overall low rates of apoptosis in this tissue may make it difficult to detect changes. Taken together, these data establish quantitative and qualitative hallmarks of follicle cell aging in the *Drosophila* ovary.

### Characterization of cell cycle regulation in aging follicle cells

The FSC lineage is highly proliferative, undergoing up to nine rounds of division before exiting the cell cycle at Stage 6 [13]. We therefore investigated whether the cell cycle of follicle cells is altered with age. First, we used an EdU assay to quantify the number of follicle cells per germarium in S-phase in ovarioles from young and aged flies (Figure 2A, A’, B, B’). Indeed, we observed a statistically significant increase in EdU^+^ cells, from 6.5 ± 1.4 cells per germarium in young flies versus 12.5 ± 2.8 cells per germarium in aged flies (Figure 2C). In contrast, we found no significant difference in the number of follicle cells that were positive for phospho-histone H3 (pHH3), which is a marker of mitosis (Figure 2D-F, D’-E’). Together, these results indicate that follicle cells in aged ovarioles are spending a greater fraction of time in S-phase but are not dividing more frequently. We hypothesized that this could be the result of follicle cells in aged germaria requiring more time to accommodate replication stress and DNA damage. Consistent with this, we found that the number of follicle cells per germarium that are positive for the DNA damage marker, gH2Av, significantly increased with age from 1.7 ± 0.03 cells per germarium to 3.4 ± 0.6 cells per germarium (Figure 2G-I).

**Figure 2:**
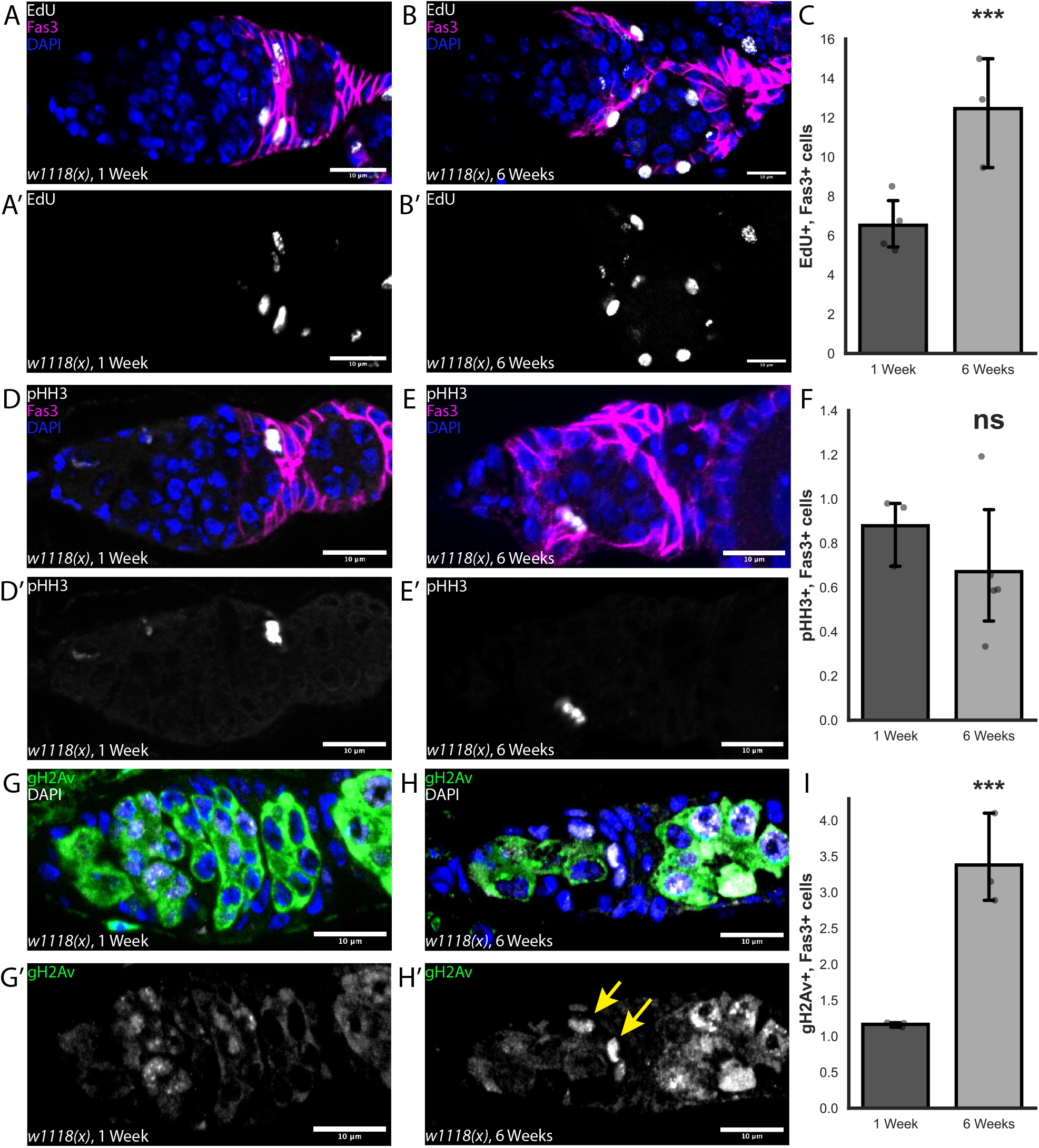
Aging leads to an extended S-phase and increased DNA damage in follicle cells. **(A-B)** Germarium from *w1118(x);;* flies at 1 (A) or 6 (B) weeks old, stained for EdU (grey), Fas3 (magenta), and DAPI (blue). **(A’-B’)** EdU identifies cells that are in or enter into S-phase during the 1-hour incubation prior to fixation. **(C)** Quantification of the number of EdU+, Fas3+ cells per germarium at 1 and 6 weeks old. n = 98 and 65 germaria, respectively. **(D-E)** Germarium from *w1118(x);;* flies at 1 (D) or 6 (E) weeks old, stained for pHH3 (grey), Fas3 (magenta), and DAPI (blue). **(D’-E’)** pHH3 identifies cells that are in mitosis. **(F)** Quantification of the number of pHH3+, Fas3+ cells per germarium at 1 and 6 weeks old. n = 200 and 124 germaria, respectively. **(G-H)** Germarium from *w1118(x);;* flies at 1 (G) or 6 (H) weeks old, stained for gH2Av (grey), Fas3 (magenta), and DAPI (blue). **(G’-H’)** gH2Av identifies sites of DNA damage. **(I)** Quantification of the number of highly positive gH2Av+, Fas3+ cells per germarium at 1 and 6 weeks old. n = 35 and 39 germaria, respectively. Scale bars are 10 µm. ns = not significant, *p<0.05, **p<0.01, ***p<0.001, using Poisson Dispersion Test (C, F, I).

### Follicle cells are unable to accommodate germ-cell cysts in aged germaria

As the appearance of severe defects in follicle formation is not associated with a substantial change in follicle cell proliferation in aged ovarioles, we next considered whether these phenotypes are associated with a loss of coordination between germ cells and follicle cells. The rate of germ-cell cyst progression from Region 2a of the germarium into the follicle epithelium in Region 2b is regulated in part by a nutrition-associated checkpoint. This checkpoint is detectable as an apoptotic cyst in Region 2a, which occurs in approximately 65% of germaria when flies are maintained on standard growth media [27]. The dying cysts can be visualized with Apoptag or TUNEL staining and are likely also the source of LysoTracker^+^ germ cells in Region 2a that were reported separately [18,27,28]. To investigate this checkpoint as a function of age, we first stained young and aged germaria with LysoTracker (Figure 3A, A’). We found that a band of LysoTracker^+^ germ cells was present in Region 2a in approximately 50% of germaria from young flies maintained under the growth conditions we used in this study. Though LysoTracker staining was incompatible with the Click-iT TUNEL assay, we found that both methods identify structures with the same frequency, shape, and location (Figure 3F-J). In addition, we found that these structures are much larger than typical lysosomes and they do not contain either the canonical lysosome marker Lamp1 or the late endosome marker Rab7 (Supplementary Figure 3A-B). Together, these observations indicate that both LysoTracker and TUNEL staining identify apoptotic cysts in Region 2a. Surprisingly, the frequency of germaria with a band of LysoTracker^+^ germ-cell cysts did not change in the aged condition (Figure 3C), indicating that this checkpoint does not decline with age. However, we found that ovarioles from aged flies without LysoTracker^+^ germ-cell cysts in Region 2a were significantly more likely to have one or more age-associated follicle cell phenotypes, whereas no such association was found in ovarioles from young flies (Figure 3B-B’, D). This suggests that the increase in follicle cell phenotypes with age is linked to a lack of coordination between germ cells and follicle cells.

**Figure 3:**
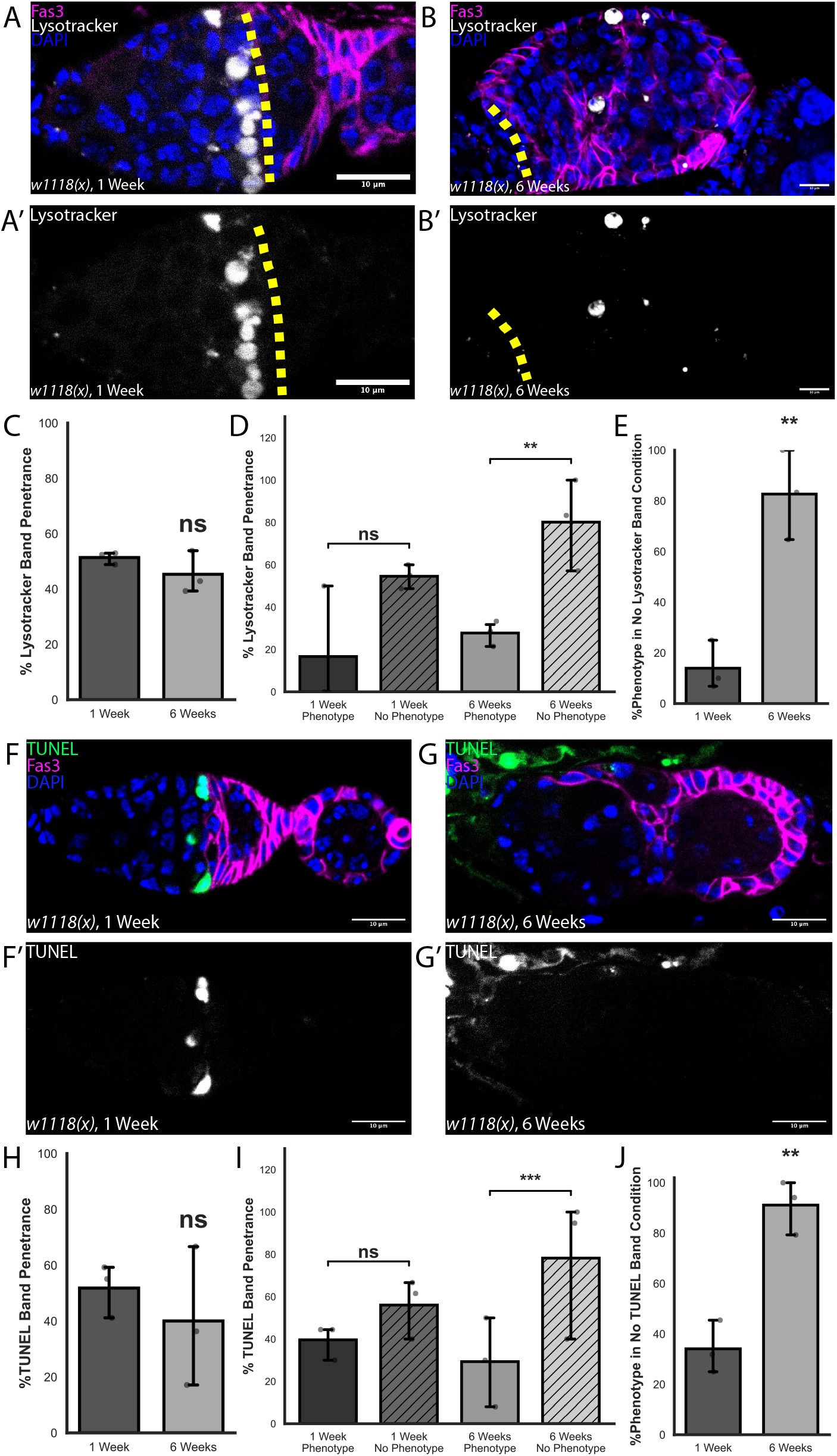
Aged ovaries cannot accommodate germ-cell cyst death. **(A-B)** Germarium from *w1118(x);;* flies at 1 (A) and 6 (B) weeks old, stained for Lysotracker (grey), Fas3 (magenta), and DAPI (blue). **(A’)** A Lysotracker-positive band may be present anterior to the Region 2a/2b border (yellow dashed line) in 1-week-old germaria. **(B’)** Lysotracker-positive bands may be absent anterior to the Region 2a/2b border (yellow dashed line) at 6-weeks old, phenotype-positive germaria. **(C)** Quantification of the percent of germaria with the Lysotracker-positive band at 1 and 6 weeks old. n = 124 and 83 germaria for 1- and 6-weeks old. **(D)** Quantification of the percent of germaria at 6 weeks old with the Lysotracker-positive band in phenotype-positive and phenotype-negative germaria. n = 57 and 26 germaria, respectively. **(E)** Quantification of phenotype penetrance in germaria in which no Lysotracker-positive band is present at 1 and 6 weeks old. n = 62 and 47, respectively. **(F-G)** Germarium from *w1118(x);;* flies at 1 (A) and 6 (B) weeks old, stained for TUNEL (grey), Fas3 (magenta), and DAPI (blue). **(F’)** A TUNEL-positive band is present anterior to the Region 2a/2b border in germaria at 1 week old. **(G’)** TUNEL-positive bands may be absent anterior to the Region 2a/2b border in phenotype-positive germaria at 6 weeks old. **(H)** Quantification of the percent of germaria with the TUNEL-positive band at 1 and 6 weeks old. n = 110 and 119 germaria, respectively. **(I)** Quantification of the percent of 6-week-old germaria with the TUNEL-positive band in phenotype-positive and phenotype-negative germaria. N = 87 and 32 germaria, respectively. **(J)** Quantification of phenotype penetrance in germaria where no TUNEL-positive band is present at 1 and 6 weeks old. n = 53 and 67 germaria, respectively. Scale bars are 10 µm. ns = not significant, *p<0.05, **p<0.01, ***p<0.001, using Welch’s t-test (C, D, H, J) and Binomial General Linearized Mixed Model (D, I).

### Single-cell RNA sequencing of aged Drosophila ovaries reveals key transcriptional differences in the follicle cell lineage

To build a comprehensive understanding of the transcriptional differences between young and aged germaria, we performed single-cell RNA sequencing (scRNA-seq) of one-week and six-week-old ovaries using 10x Genomics system and lllumina sequencing. This produced transcriptional data for 20,818 cells across two timepoints (Supplementary Table 1) after applying standard quality control filters (Supplementary Figure 4A-B). Using the standard Seurat workflow, we clustered the cells into nine broad clusters (Figure 4A): Germ cells (*vas*+), Escort cells (*Wnt4*+, *Wnt6*+), FSCs + pFCs (*zfh1*+, *cas*+, *eya*+, *Fas3*+), Early-Mid FCs (*eya*+, *br*+, *Fas3*+, *mid*+), Late FCs (*br*+, *mid*+, *Yp1*+), Muscle (*Mhc*+), Immune (*atilla*+, *nec*+), and Spermatheca (*lz*+), and Neurons (*pros*+, *nrv3*+) (Supplementary Figure 5A) [10,29–33]. We mapped the cells from aged ovaries onto the young ovary dataset using the Seurat integration features (Figure 4B). Mean silhouette scores revealed that most ovarian cell types maintained strong transcriptional separability across age, indicating strong clustering identities (Supplementary Figure 5B). Notably, we found that the Early-Mid FCs have the highest number of differentially expressed genes, followed by FSCs + pFCs and muscle, with germ cells ranking eighth (Figure 4C). In looking at the whole aged ovary, we found 321 differentially expressed genes (DEGs) between the young and aged condition, and 156 DEGs specifically in the FSCs + pFCs cluster (Figure 4D, Supplementary Figure 5C, Supplementary Tables 4-5). To determine how these transcriptional changes relate to the transcriptional changes in mammalian granulosa cells with age, we merged the FSCs + pFCs cluster with the Early-Mid FCs cluster to create a more “granulosa-like” cell cluster, identified the mammalian orthologs of the DEGs in this merged cluster and cross-referenced them with a list of age-associated DEGs from a recently published study of mouse and human aged granulosa cells [19]. Interestingly, we found several genes that are significantly upregulated in both mouse and human granulosa cells that have homologous genes that are significantly upregulated in our aged *Drosophila* “granulosa-like” cell cluster (Figure 4F). This list includes genes involved in signal transduction, such as the PDGF-related factor *Pvf1*, and the JNK-stress pathway effector *Jun-related antigen*; and immune response genes such as the NFKB1 homolog, *Relish*, and an uncharacterized serine hydrolase, CG10472, that is homologous to mammalian CTRB1 and is predicted to be involved in innate immunity.

**Figure 4:**
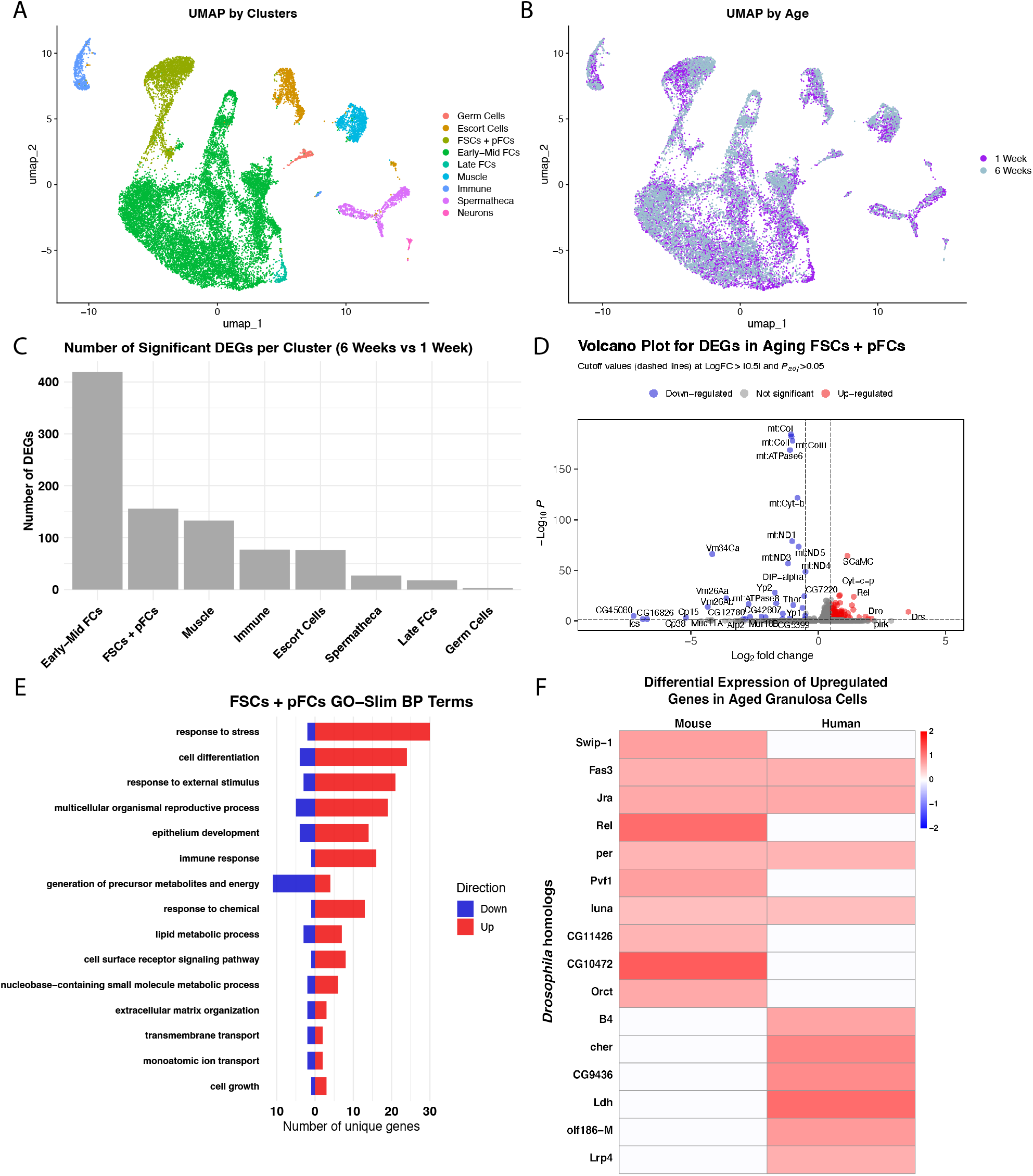
Single-cell sequencing of young and aged *Drosophila* ovaries reveals large transcriptional differences in aged follicle cells. **(A)** UMAP of the merged dataset, annotated by the 8 clusters. **(B)** UMAP of the merged dataset, annotated by the 1 and 6-week old datasets. **(C)** Ranking of the number of significant differentially expressed genes (DEGs) in each cluster, comparing the 6-week to the 1-week old datasets. **(D)** Volcano plot of DEGs from the FSCs + pFCs cluster. Blue dots are down-regulated genes, grey dots are not significant genes, and red dots are up-regulated genes. Significance is indicated by an absolute value log fold-change greater than 0.5 and adjusted p-value greater than 0.05. **(E)** Gene ontology terms, using GO-Slim for the FSCs + pFCs cluster. **(F)** Heatmap of fly genes with mammalian homologs that are up-regulated with age, from Gaylord et al, 2025 [19].

### Overexpression of Atg8a in early follicle cells suppresses age-related follicle cell decline and rescues reproductive capacity

To identify genetic modifications that could counteract the age-associated decline in fertility, we screened through well-known genes associated with classic hallmarks of aging, such as cell differentiation, response to stress, and epithelium development (Figure 4E, Supplementary Figure 5D). Interestingly, we noticed that overexpression of Atg8a::mCherry-GFP specifically in the early FSC lineage using *109-30-Gal4* produced particularly healthy flies. *Atg8a* is an autophagy gene that is essential for the initiation of an autophagosome and fusion of the autophagosome to a lysosome [34–36] and, while it did not come up as a significant DEG in our single-cell RNA sequencing dataset (Supplementary Figure 5E), autophagy defects are highly associated with age-associated defects. We therefore investigated whether upregulating *Atg8a* in follicle cells could improve the age-related decline in ovarian function.

Indeed, we found that there was a significant reduction in the frequency of follicle cell phenotypes relative to an internal control in aged ovaries (Figure 5A-C). Interestingly, we observed a slight increase in follicle cell phenotype frequency in young ovaries (Figure 5C), suggesting that at least some of the beneficial effects of *Atg8a* overexpression may be specific to the aged condition. In addition, we observed a significant increase in the percentage of aged germaria that have maintained the ring of N-cad upon overexpression of Atg8a::mCherry-GFP compared to control (Figure 1G’, H’). Moreover, we found that overexpression of Atg8a::mCherry-GFP decreased the number of highly expressing gH2Av positive follicle cells (Figure G-H, G’-H’, I). Additionally, we found that overexpression of Atg8::mCherry-GFP in follicle cells has cell non-autonomous effects on the aged germline. While overexpression of Atg8::mCherry-GFP did not significantly alter the frequency of germaria with LysoTracker^+^ germ-cell cysts in Region 2a, it eliminated the association between the presence of follicle cell phenotypes and the lack a LysoTracker^+^ cyst in Region 2a (Figure 6A-D). Finally, we assayed for egg-laying to determine if overexpression of *Atg8a* in follicle cells could rescue age-related decline in reproductive capacity [38]. We found that egg-laying increased from 2 ± 1.5 eggs per female per day in aged control *Drosophila* to 5.9 ± 2.6 eggs per female per day in *Atg8a* overexpression condition, a nearly 2-fold increase in egg-laying (Figure 6F). Importantly, egg-laying in aged *Atg8a* overexpression condition was not significantly different from that of young control *Drosophila* (Figure 6F). Taken together, these data indicate that overexpression of *Atg8a* improves age-related follicle cell decline in tissue integrity, cell adhesion, DNA damage repair, and reproductive capacity in aged ovaries.

**Figure 5:**
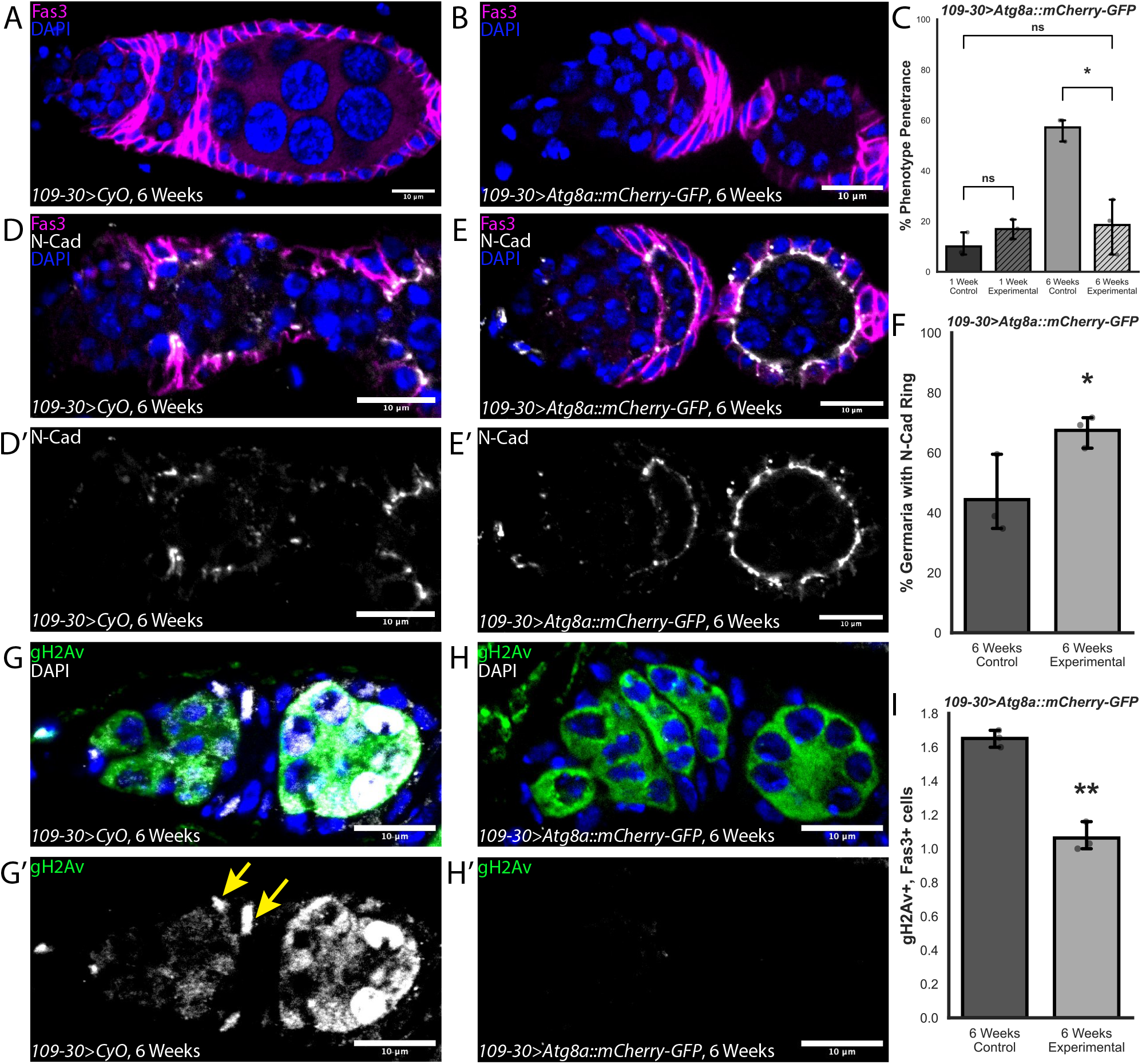
Overexpression of *Atg8a* in follicle cells suppresses age-related follicle cell decline. **(A)** Germarium from a *109-30>CyO* control fly at 6 weeks old stained for Fas3 (magenta) and DAPI (blue). **(B)** Germarium from a*109-30;>;Atg8a::mCherry-GFP;* experimental fly stained for Fas3 (magenta) and DAPI (blue). **(C)** Quantification of percent of germaria with follicle cell phenotypes in 1- and 6-week-old *;109-30;>;CyO;* and *;109-30;>;Atg8a::mCherry-GFP;*. n = 115 and 235 germaria for 1-week-old *;109-30;>;CyO;* and *;109-30;>;Atg8a::mCherry-GFP;*, and n = 61 and 140 for 6-weeks-old *;109-30;>;CyO;* and *;109-30;>;Atg8a::mCherry-GFP;*. **(D)** Germarium from a 6-week-old, *;109-30;>;CyO;* fly, stained for N-cad (grey), Fas3 (magenta), and DAPI (blue). **(D’)** Loss of N-cadherin ring around developing germ-cell cysts in 6-weeks-old *;109-30;>;CyO;*. **(E)** Germarium from a 6-week-old *;109-30;>;Atg8a::mCherry-GFP;* fly, stained for N-cad (grey), Fas3 (magenta), and DAPI (blue). **(E’)** Rescue of N-cadherin ring around developing germ-cell cysts in 6-week-old *;109-30;>;Atg8a::mCherry-GFP;* germaria. **(F)** Quantification of the percent of germaria with the N-cadherin ring around germ-cell cysts in 6-week-old *;109-30;>;CyO; and ;109-30;>;Atg8a::mCherry-GFP;*. n = 101 and 118 for 6-weeks-old *;109-30;>;CyO;* and *;109-30;>;Atg8a::mCherry-GFP;*. **(G)** Germarium from a 6-week-old *;109-30;>;CyO;* control fly stained for gH2Av (grey), Fas3 (magenta), and DAPI (blue). **(H)** Germarium from a 6-week-old *;109-30;>;Atg8a::mCherry-GFP;* control fly stained for gH2Av (grey), Fas3 (magenta), and DAPI (blue). **(G’-H’)** gH2Av identifies sites of DNA damage. **(I)** Quantification of the number of highly positive gH2Av+, Fas3+ cells per germarium in 6-week-old *;109-30;>;CyO; and ;109-30;>;Atg8a::mCherry-GFP;*. n = 77 and 84 for 6-weeks-old *;109-30;>;CyO;* and *;109-30;>;Atg8a::mCherry-GFP;*. Scale bars are 10 µm. ns = not significant, *p<0.05, **p<0.01, ***p<0.001, using Welch’s t-test (C), Student’s t-test (F), and Poisson Dispersion Test (I).

**Figure 6.**
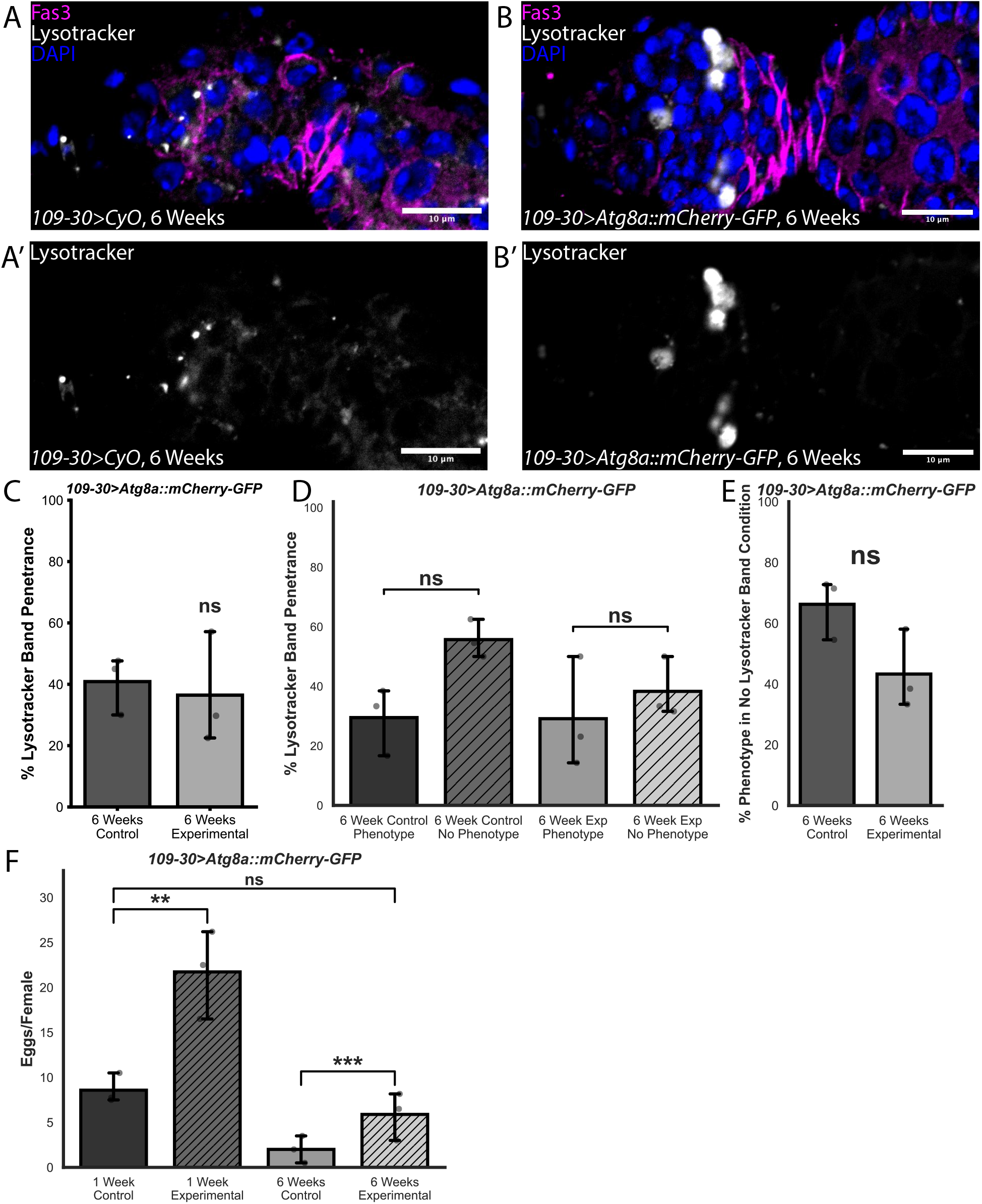
Overexpression of *Atg8a* in follicle cells has non-autonomous effect on germ-cell cyst regulation and rescues egg laying. **(A)** Germarium from a 6-week-old *;109-30;>;CyO;* control fly, stained for Lysotracker (grey), Fas3 (magenta), and DAPI (blue). **(A’)** A Lysotracker-positive band may be absent anterior to the Region 2a/2b border) in phenotype-positive germaria. **(B)** Germarium from a 6-week-old *;109-30;>;Atg8a::mCherry-GFP;* experimental fly, stained for Lysotracker (grey), Fas3 (magenta), and DAPI (blue). **(B’)** Lysotracker-positive bands may be present anterior to the Region 2a/2b border in phenotype-negative germaria. **(C)** Quantification of the percent of germaria with the Lysotracker-positive band in 6-week-old *;109-30;>;CyO; and ;109-30;>;Atg8a::mCherry-GFP;*. n = 51 and 84 germaria for *;109-30;>;CyO;* and *;109-30;>;Atg8a::mCherry-GFP*. **(D)** Quantification of the percent of 6-week-old *;109-30;>;CyO;* germaria and percent of 6-week-old *;109-30;>;Atg8a::mCherry-GFP;* with the Lysotracker-positive band in phenotype-positive and phenotype-negative germaria. n = 28 *;109-30;>;CyO;* phenotype-positive germaria, n = 23 *;109-30;>;CyO;* phenotype-negative germaria, n = 36 *;109-30;>;Atg8a::mCherry-GFP;* phenotype-positive germaria, n = 49 *;109-30;>;Atg8a::mCherry-GFP;* phenotype-negative germaria. **(E)** Quantification of phenotype penetrance in germaria where no Lysotracker-positive band is present for *;109-30;>;CyO;* and *;109-30;>;Atg8a::mCherry-GFP*. n = 29 and 60 germaria for *;109-30;>;CyO;* and *;109-30;>;Atg8a::mCherry-GFP*. **(F)** Quantification of eggs laid per female in a 24-hour period for 1- and 6-week-old *;109-30;>;CyO;* and *;109-30;>;Atg8a::mCherry-GFP* flies. n = 10 and 11 vials for 1-week-old *;109-30;>;CyO;* and *;109-30;>;Atg8a::mCherry-GFP;*, and n = 5 and 8 vials for 6-weeks-old *;109-30;>;CyO;* and *;109-30;>;Atg8a::mCherry-GFP;*. Scale bars are 10 µm. ns = not significant, *p<0.05, **p<0.01, ***p<0.001, using Welch’s t-test (C, E), Binomial General Linearized Mixed Model (D), and Poisson Distribution Test (F).

## Discussion

Here, we have identified several key genetic and cellular changes that accumulate with age in the *Drosophila* follicle epithelium and discovered a targeted intervention, overexpression of *Atg8a* specifically in the early FSC lineage, that ameliorates these phenotypes. Several of the defects we identified, such as gaps in the follicle epithelium and the lack of an interfollicular stalk, are incompatible with follicle development and thus likely to be significant causes of reduced fecundity in aged flies. In addition, we identified intracellular phenotypes, including increased DNA damage and a disruption of cellular junctions, that indicate a loss of normal follicle cell function. Moreover, our scRNAseq data provide a comprehensive account of cell type specific changes in gene expression with age. Interestingly, our analysis predicts that there are more differentially expressed genes in the follicle lineage than any other cell type in the ovary. Additionally, several of the highly upregulated genes in mammalian aged granulosa cells are homologous to significantly upregulated genes in the aged *Drosophila* follicle lineage. Taken together, these observations emphasize the importance of the early FSC lineage in *Drosophila* ovarian aging and provide a baseline description of age-associated changes in this tissue that can be used to evaluate the effects of experimental interventions that target follicle cells. Notably, the follicle cell phenotypes we observed, including DNA damage and cell cycle alterations, are also hallmarks of reproductive aging in mammals [39], highlighting the opportunity to understand conserved mechanisms of reproductive aging through these studies.

We were surprised to find that overexpression of *Atg8a* with *190-30-Gal4* is sufficient to increase fecundity in both young and old flies. *109-30-Gal4* expression is restricted to a relatively small population of somatic cells located in the posterior half of the germarium and the first 2-3 budded follicles, yet genetic manipulation of this small population increased the overall output of the whole ovary. This demonstrates that oogenesis is not operating at maximum capacity under the growth conditions we used, and that the function of these cells in the early FSC lineage is rate-limiting. In addition, though fecundity still declines with age in flies with *Atg8a* overexpression, it is notable that the rate of egg laying in the aged condition is not significantly different from that of young control flies. Thus, despite the impact of other sources of decline that inevitably occur with age, these flies still maintain a youthful level of fecundity in the aged condition. These findings are consistent with emerging evidence that targeting the effects of ovarian somatic cell aging in mammals may slow reproductive aging. For example, the pro-inflammatory signal, CD38, which is primarily expressed in endothelial and immune cells, increases with age and global inhibition of this increase enhances fertility in aged mice [39]. Likewise, interventions that reduce fibrosis caused by ovarian somatic cells are being considered as possible therapeutic avenues [41]. An advantage of these interventions is that, by focusing on the support cells, they avoid direct manipulation of the germ cells that will contribute to the next generation. The genetic tools available here enabled us to use targeted genetic manipulations to study the role of somatic cells in reproductive aging.

The cells of the early FSC lineage may influence the rate of oogenesis through their direct role in facilitating follicle formation as well as through signaling with germ cells that helps coordinate the supply of follicle cells with demand from newly-produced germ-cell cysts. A key factor in this coordination process is the nutrition-dependent checkpoint at Region 2a. This checkpoint and a second one that occurs in mid oogenesis may be orthologous to a process in the mammalian ovary in which granulosa cells secrete signals to induce follicle atresia [42]. We found that the frequency of germaria with a dying cyst in Region 2a did not change between young and old flies or even with the overexpression of *Atg8*. However, the presence of age-associated follicle cell phenotypes is linked to the absence of a dying cyst in Region 2a, raising the interesting possibility that germaria without a recent activation of this checkpoint are more susceptible to a loss of coordination between germ cells and follicle cells. Alternatively, germaria with follicle cell phenotypes may be less likely to activate the checkpoint, possibly leading to further loss of coordination. Understanding how a loss of coordination at this stage of oogenesis contributes to overall fecundity rates will be an important topic for future studies.

Mechanistically, *Atg8a*, which is homologous to mammalian LC3, is well-studied as a key regulator of autophagy [43,44]. Autophagy is a protective cellular function that promotes the degradation of proteins, organelles, and other cellular components, to mitigate the accumulation of damaged biomolecules over time [44]. Autophagy is regulated by a cascade of signals, with several ATG (autophagy-related genes) involved [36,44]. *Atg8a* resides on the membrane of autophagosomes during formation and expansion and is known to increase in expression in response to stress conditions such as starvation and reactive oxygen species accumulation in the *Drosophila* ovary [34–36]. In addition, an upstream regulator, *Atg7*, is required in follicle cells for coordination between germ cells and follicle cells in the progression of oogenesis [35]. In the mammalian ovary, LC3 expression and standard indicators of autophagy activity, such as lysosome number and autolysosome fusion, decline with age, thus reducing the capacity of this protective mechanism [44,45]. Autophagy is critical for DNA damage mediation, where autophagy is promoted by the DNA damage response and repair pathways [47,48]. Additionally autophagy, and Atg8a specifically, also works upstream to prevent DNA damage and oxidative stress [49,50]. Consistent with this, we found that overexpression of *Atg8a* decreases the accumulation of DNA damage in follicle cells. Likewise, autophagy machinery is known to traffic E-cadherin to lysosomes [51]. This suggests that similar mechanisms may facilitate the maintenance of proper N-cad localization in ovarioles with *Atg8a* overexpression, as we have shown. These anti-aging effects at the cellular level likely contribute to the improved function of the follicle epithelium at the tissue level.

In summary, this study contributes to a growing body of literature that highlights the importance of ovarian somatic cells in reproductive health and aging. Further studies into other features of follicle cell aging and how they contribute to the decline in overall ovarian function will be of interest. In addition, with emerging evidence that ovarian health contributes to overall organism health, our findings will support future efforts to investigate the effects of follicle cell health on other organ systems and lifespan.

## Supporting information

Cluster Counts

Top 50 DEGs by cluster

DEG counts by cluster

List of whole ovary DEGs

List of FSCs + pFCs cluster DEGs

## Acknowledgements

We are grateful to Stephane Noselli and Gábor Juhász for sharing reagents and fly stocks. We thank Vanessa Huynh for technical support, Katja Rust for the schematics in Figure 1A-B, Nathaniel Paul Meyer for help with the single-cell RNA sequencing, and Eliza Gaylord and Ryan Samuel for the mouse and human DEG lists. We also thank Diana Laird for feedback on this manuscript. We thank the Bloomington Drosophila Stock Center (BDSC) (NIH P40OD018537) for stocks obtained used in this study. We thank the Developmental Studies Hybridoma Bank (DSHB), created by the NICHD of the NIH and maintained at The University of Iowa, for antibodies used in this study. This study was supported by the National Institute of General Medical Sciences of the National Institutes of Health under Award Number R35GM136348 to T.G.N, a Bakar Aging Research Institute Investigator Award to T.G.N., the Eunice Kennedy Shriver National Institute Of Child Health & Human Development of the National Institutes of Health under Award Number T32HD007470, and a National Science Foundation Graduate Research Fellowship under Grant No 2445150 to E.A.W..

## Figure Legends

**Supplementary Figure 1.**
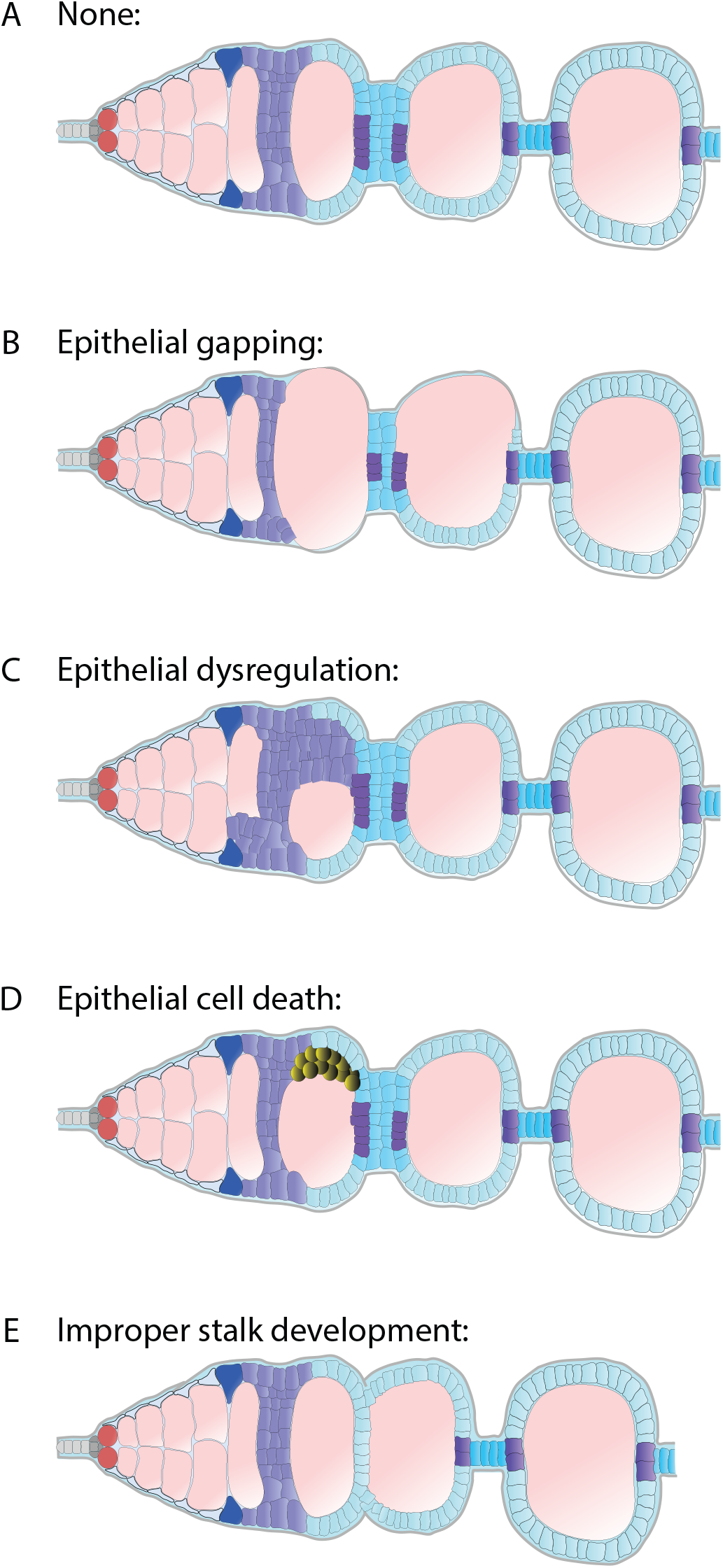
Schematics of aging phenotypes for characterizations. **(A)** Schematic of no phenotype condition. **(B)** Schematic of epithelial gapping phenotype, where germ cysts are not fully encapsulated by early follicle cells. **(C)** Schematic of dysregulated epithelium phenotype, where early follicle cells are not forming a single-layer epithelium. **(D)** Schematic of epithelial cell death phenotype, where large groups of follicle cells appear rounded, rather than cuboidal. **(E)** Schematic of an example of impaired stalk development phenotype, where stalk cells may not be present between developing follicles.

**Supplementary Figure 2.**
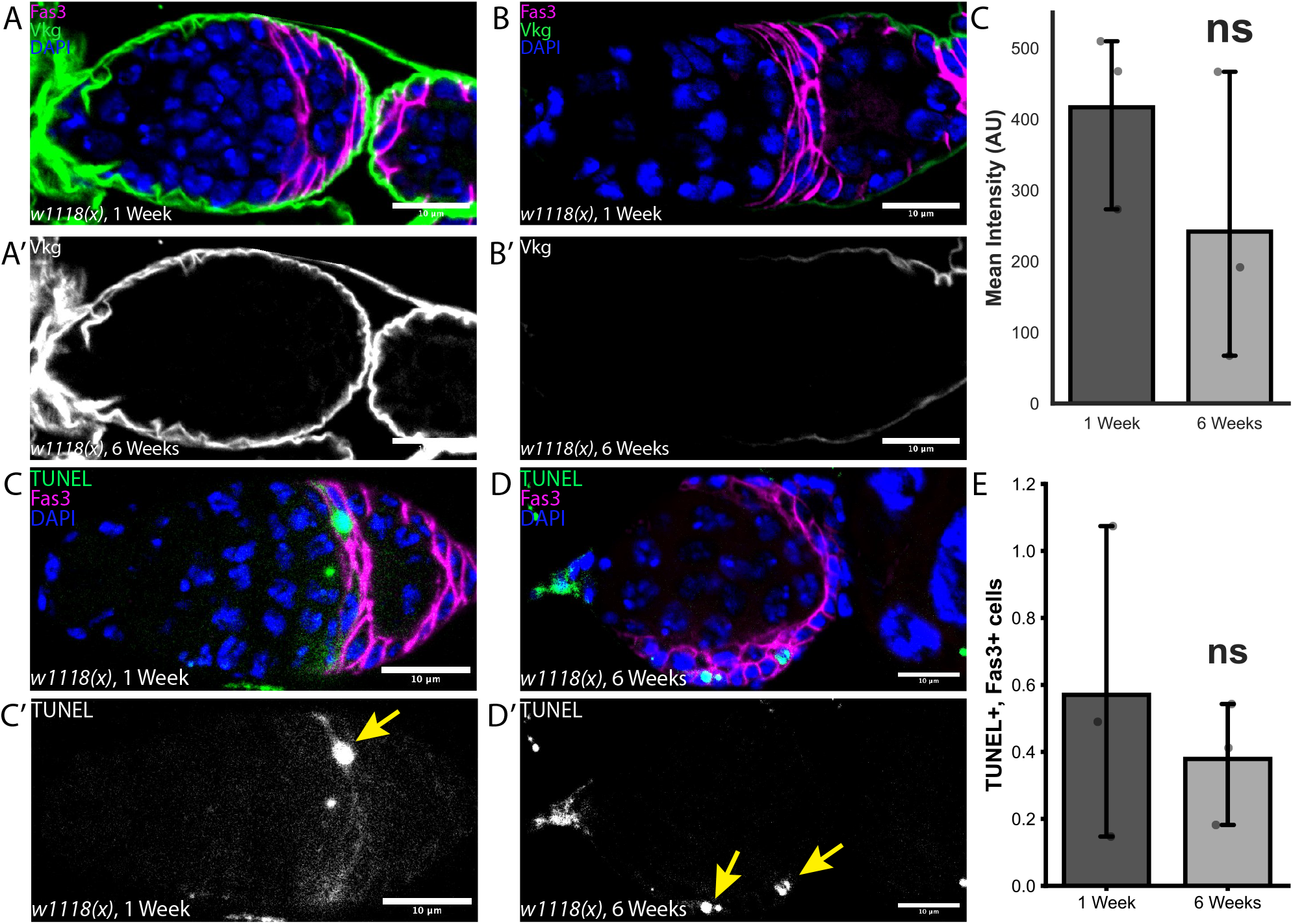
Further characteristics of the aging ovary. **(A-B)** Germarium from a 1-week-old (A) and 6-week-old (B) *w1118(x);;* flies, stained for Vkg (green), Fas3 (magenta), and DAPI (blue). **(A’-B’)** Vkg stains for collagen IV, which adheres to the perimeter of the germarium. **(C)** Quantification of the average intensity of Vkg around the perimeter of the germarium at 1- and 6-weeks-old. n = 26 and 27 germaria for 1- and 6-weeks-old. **(D-E)** Germarium from a 1-week-old (C) and 6-week-old (D) *w1118(x);;* flies, stained for TUNEL (green), Fas3 (magenta), and DAPI (blue). **(D’-E’)** TUNEL stains sites of DNA fragmentation, indicating final stages of apoptosis. **(F)** Quantification of the average number of TUNEL-positive, Fas3-positive cells per germarium at 1- and 6-weeks-old. n = 110 and 119 germaria for 1- and 6-weeks old. Scale bars are 10 µm. ns = not significant, *p<0.05, **p<0.01, ***p<0.001, using Welch’s t-test (C) and Poisson Distribution Test (F).

**Supplementary Figure 3.**
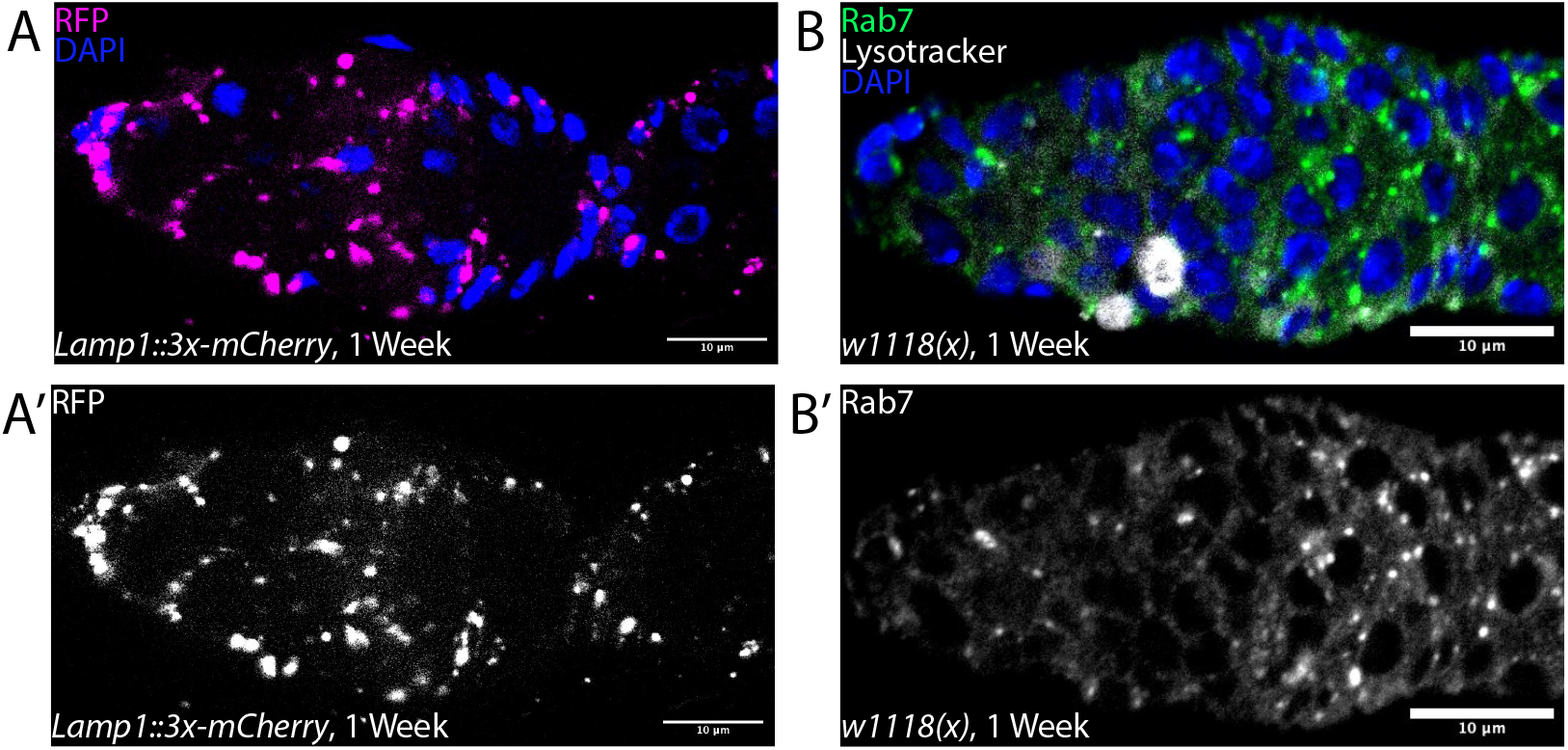
Large Lysotracker-positive structures in Region 2a are not canonical lysosomes or late endosomes. **(A)** Germarium from a 1-week-old *;;Lamp1::3x-mCherry* fly, stained for RFP (magenta) and DAPI (blue). **(A’)** Lamp1 is a membrane protein on lysosomes and does not have the same pattern as the large Lysotracker-positive structures seen in Region 2a. **(B)** Germarium from a 1-week-old *;;w1118(x)* fly, stained for Rab7 (green), Lysotracker (grey), and DAPI (blue). **(B’)** Rab7 is a protein associated with late endosomes and does not colocalize with the large Lysotracker-positive structures seen in Region 2a.

**Supplementary Figure 4.**
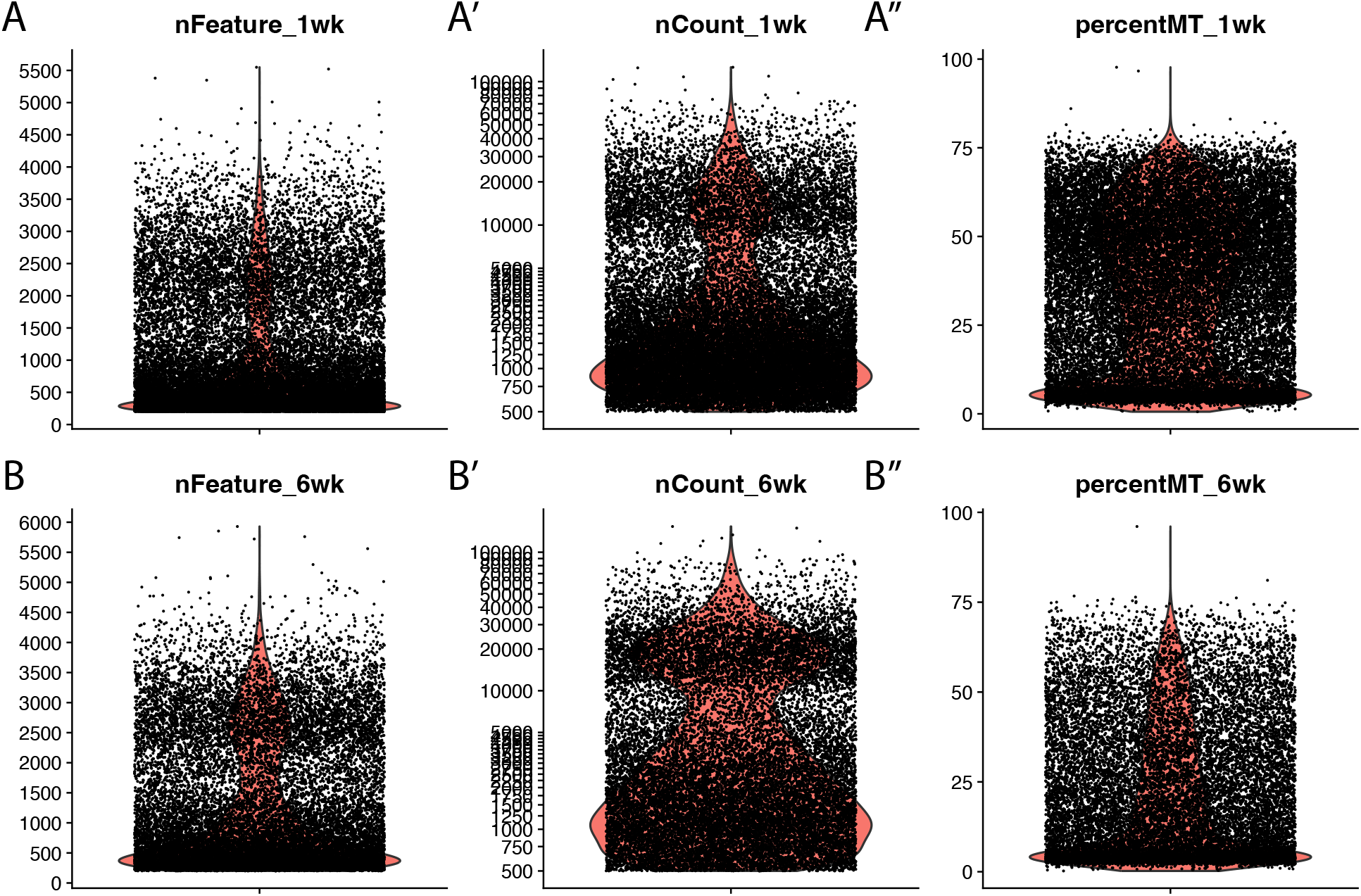
Quality control plots for single-cell RNA sequencing of young and aged ovaries. **(A-B)** Violin plots for the number of RNA features by age and replicate. **(A’-B’)** Violin plots for the number of RNA counts by age and replicate. **(A’’-B’’)** Violin plots for the percent mitochondria reads per cell ID by age and replicate.

**Supplementary Figure 5.**
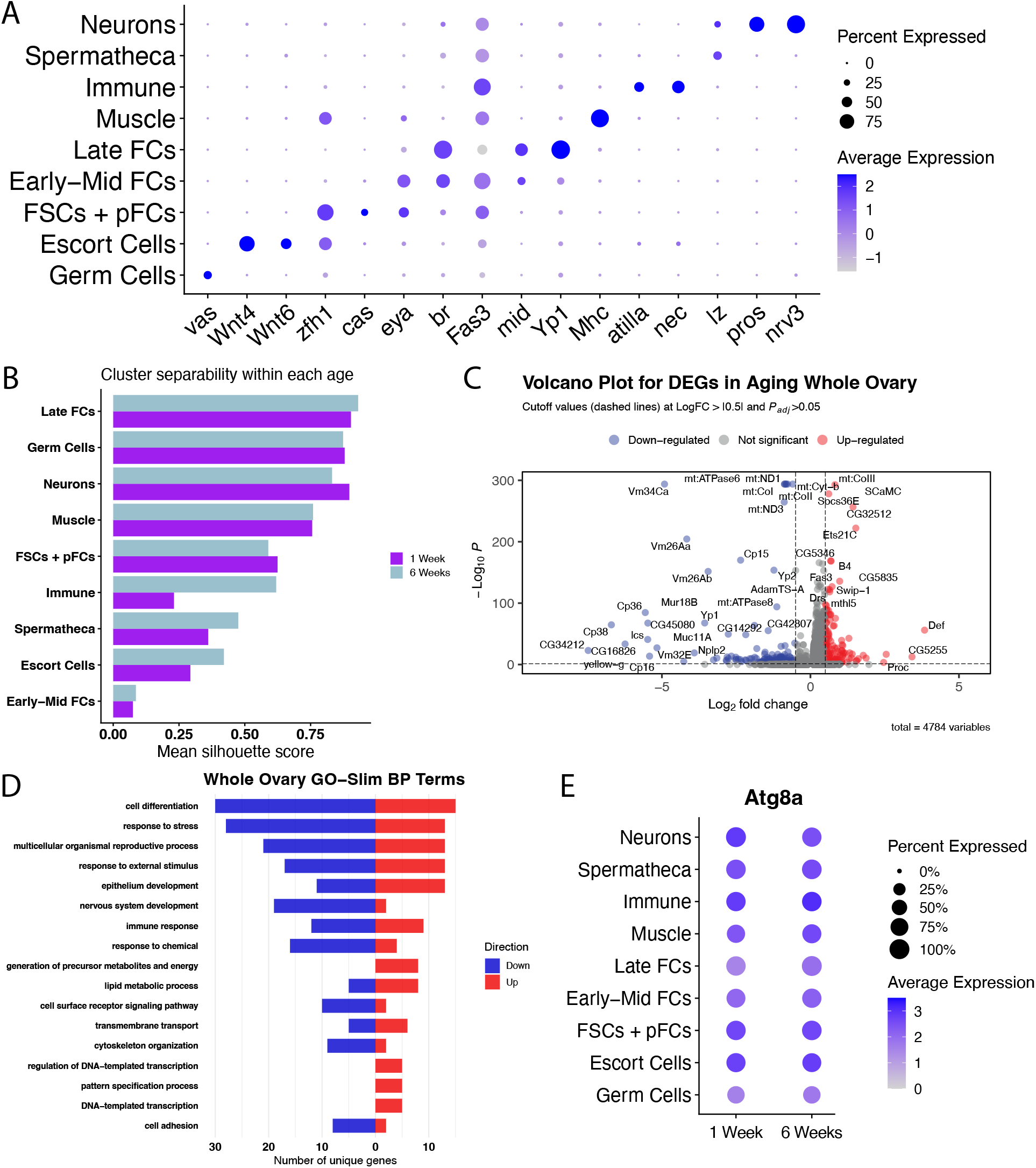
Cluster identification and additional analysis of single-cell RNA sequencing of young and aged ovaries. **(A**) Dot plot of gene expression for markers associated with each cluster of cells. **(B)** Mean silhouette score for each cluster compared against its own dataset for 1- and 6-weeks-old. **(C)** Volcano plot of DEGs for the whole ovary. Blue dots are down-regulated genes, grey dots are not significant genes, and red dots are up-regulated genes. Significance is indicated by an absolute value log fold-change greater than 0.5 and adjusted p-value greater than 0.05. **(D)** Gene ontology terms, using GO-Slim for the whole ovary. **(E)** Dot plot of gene expression for *Atg8a* in each cluster for 1- and 6-weeks-old.

**Supplementary Table 1. Cell counts by dataset**.

**Supplementary Table 2. Top 50 DEGs by cluster comparing the 1- and 6-week-old datasets**.

**Supplementary Table 3. Number of significant DEGs by cluster comparing the datasets from 1 and 6 weeks old**.

**Supplementary Table 4. List of the 321 significant DEGs for whole aging ovary. Supplementary Table 5. List of the 156 significant DEGs for whole aging ovary**.

## References

1. López-Otín C, Blasco MA, Partridge L, Serrano M, Kroemer G. The hallmarks of aging. Cell. 2013;153: 1194–1217.

2. Desai S, Rajkovic A. Genetics of reproductive aging from gonadal dysgenesis through menopause. Semin Reprod Med. 2017;35: 147–159.

3. Spradling AC, Niu W, Yin Q, Pathak M, Maurya B. Conservation of oocyte development in germline cysts from Drosophila to mouse. Elife. 2022;11. doi:10.7554/eLife.83230

4. Fellmeth JE, McKim KS. A brief history of Drosophila (female) meiosis. Genes (Basel). 2022;13: 775.

5. Doherty CA, Amargant F, Shvartsman SY, Duncan FE, Gavis ER. Bidirectional communication in oogenesis: a dynamic conversation in mice and Drosophila. Trends Cell Biol. 2022;32: 311–323.

6. Stebbings LA, Todman MG, Phillips R, Greer CE, Tam J, Phelan P, et al. Gap junctions in Drosophila: developmental expression of the entire innexin gene family. Mech Dev. 2002;113: 197–205.

7. Carabatsos MJ, Sellitto C, Goodenough DA, Albertini DF. Oocyte-granulosa cell heterologous gap junctions are required for the coordination of nuclear and cytoplasmic meiotic competence. Dev Biol. 2000;226: 167–179.

8. Belles X, Piulachs M-D. Ecdysone signalling and ovarian development in insects: from stem cells to ovarian follicle formation. Biochim Biophys Acta. 2014. doi:10.1016/j.bbagrm.2014.05.025

9. Krajnik K, Mietkiewska K, Skowronska A, Kordowitzki P, Skowronski MT. Oogenesis in women: From molecular regulatory pathways and maternal age to stem cells. Int J Mol Sci. 2023;24. doi:10.3390/ijms24076837

10. Rust K, Nystul T. Signal transduction in the early Drosophila follicle stem cell lineage. Curr Opin Insect Sci. 2020;37: 39–48.

11. Sahai-Hernandez P, Castanieto A, Nystul TG. Drosophila models of epithelial stem cells and their niches: Drosophilamodels of epithelial stem cells. Wiley Interdiscip Rev Dev Biol. 2012;1: 447–457.

12. King RC, Rubinson AC, Smith RF. Oogenesis in adult Drosophila melanogaster. Growth. 1956;20: 121–157.

13. Margolis J, Spradling A. Identification and behavior of epithelial stem cells in the Drosophila ovary. Development. 1995;121: 3797–3807.

14. Piper MDW, Partridge L. Drosophila as a model for ageing. Biochim Biophys Acta Mol Basis Dis. 2018;1864: 2707–2717.

15. Novoseltsev VN, Arking R, Novoseltseva JA, Yashin AI. Evolutionary optimality applied to Drosophila experiments: hypothesis of constrained reproductive efficiency. Evolution. 2002;56: 1136–1149.

16. Jones DL. Aging and the germ line: where mortality and immortality meet. Stem Cell Rev. 2007;3: 192–200.

17. Grandison RC, Piper MDW, Partridge L. Amino-acid imbalance explains extension of lifespan by dietary restriction in Drosophila. Nature. 2009;462: 1061–1064.

18. Singh T, Lee EH, Hartman TR, Ruiz-Whalen DM, O’Reilly AM. Opposing Action of Hedgehog and Insulin Signaling Balances Proliferation and Autophagy to Determine Follicle Stem Cell Lifespan. Dev Cell. 2018;46: 720-734.e6.

19. Gaylord EA, Foecke MH, Samuel RM, Soygur B, Detweiler AM, McIntyre TI, et al. Comparative analysis of human and mouse ovaries across age. Science. 2025;390: eadx0659.

20. Wang H, Huang Z, Shen X, Lee Y, Song X, Shu C, et al. Rejuvenation of aged oocyte through exposure to young follicular microenvironment. Nat Aging. 2024;4: 1194–1210.

21. Fadiga J, Nystul TG. The follicle epithelium in the Drosophila ovary is maintained by a small number of stem cells. Elife. 2019;8. doi:10.7554/eLife.49050

22. Simon AF, Shih C, Mack A, Benzer S. Steroid control of longevity in Drosophila melanogaster. Science. 2003;299: 1407–1410.

23. Hegedűs K, Takáts S, Boda A, Jipa A, Nagy P, Varga K, et al. The Ccz1-Mon1-Rab7 module and Rab5 control distinct steps of autophagy. Mol Biol Cell. 2016;27: 3132–3142.

24. Meyer N, Peralta J, Nystul T. Preparation of Drosophila ovarioles for single-cell RNA sequencing. Methods Mol Biol. 2023;2626: 323–333.

25. Yonekura S, Ting C-Y, Neves G, Hung K, Hsu S-N, Chiba A, et al. The variable transmembrane domain of Drosophila N-cadherin regulates adhesive activity. Mol Cell Biol. 2006;26: 6598–6608.

26. Crowley LC, Marfell BJ, Waterhouse NJ. Detection of DNA fragmentation in apoptotic cells by TUNEL. Cold Spring Harb Protoc. 2016;2016: db.prot087221.

27. Drummond-Barbosa D, Spradling AC. Stem cells and their progeny respond to nutritional changes during Drosophila oogenesis. Dev Biol. 2001;231: 265–278.

28. Nezis IP, Lamark T, Velentzas AD, Rusten TE, Bjørkøy G, Johansen T, et al. Cell death during Drosophila melanogaster early oogenesis is mediated through autophagy. Autophagy. 2009;5: 298–302.

29. Sun J, Spradling AC. Ovulation in Drosophila is controlled by secretory cells of the female reproductive tract. Elife. 2013;2: e00415.

30. Honti V, Kurucz E, Csordás G, Laurinyecz B, Márkus R, Andó I. In vivo detection of lamellocytes in Drosophila melanogaster. Immunol Lett. 2009;126: 83–84.

31. Robertson AS, Belorgey D, Lilley KS, Lomas DA, Gubb D, Dafforn TR. Characterization of the necrotic protein that regulates the Toll-mediated immune response in Drosophila. J Biol Chem. 2003;278: 6175–6180.

32. Palmateer CM, Artikis C, Brovero SG, Friedman B, Gresham A, Arbeitman MN. Single-cell transcriptome profiles of Drosophila fruitless-expressing neurons from both sexes. Elife. 2023;12. doi:10.7554/eLife.78511

33. Roy M, Sivan-Loukianova E, Eberl DF. Cell-type-specific roles of Na+/K+ ATPase subunits in Drosophila auditory mechanosensation. Proc Natl Acad Sci U S A. 2013;110: 181–186.

34. Xie Z, Nair U, Klionsky DJ. Atg8 controls phagophore expansion during autophagosome formation. Mol Biol Cell. 2008;19: 3290–3298.

35. Barth JMI, Szabad J, Hafen E, Köhler K. Autophagy in Drosophila ovaries is induced by starvation and is required for oogenesis. Cell Death Differ. 2011;18: 915–924.

36. Bali A, Shravage BV. Characterization of the Autophagy related gene-8a (Atg8a) promoter in Drosophila melanogaster. Int J Dev Biol. 2017;61: 551–555.

37. Partridge L, Fowler K, Trevitt S, Sharp W. An examination of the effects of males on the survival and egg-production rates of female Drosophila melanogaster. J Insect Physiol. 1986;32: 925–929.

38. Balough JL, Dipali SS, Velez K, Kumar TR, Duncan FE. Hallmarks of female reproductive aging in physiologic aging mice. Nat Aging. 2024;4: 1711–1730.

39. Yang Q, Chen W, Cong L, Wang M, Li H, Wang H, et al. NADase CD38 is a key determinant of ovarian aging. Nat Aging. 2024;4: 110–128.

40. Umehara T, Winstanley YE, Andreas E, Morimoto A, Williams EJ, Smith KM, et al. Female reproductive life span is extended by targeted removal of fibrotic collagen from the mouse ovary. Sci Adv. 2022;8: eabn4564.

41. Pritchett TL, Tanner EA, McCall K. Cracking open cell death in the Drosophila ovary. Apoptosis. 2009;14: 969–979.

42. Kabeya Y, Mizushima N, Ueno T, Yamamoto A, Kirisako T, Noda T, et al. LC3, a mammalian homologue of yeast Apg8p, is localized in autophagosome membranes after processing. EMBO J. 2000;19: 5720–5728.

43. Gatica D, Lahiri V, Klionsky DJ. Cargo recognition and degradation by selective autophagy. Nat Cell Biol. 2018;20: 233–242.

44. Peters AE, Caban SJ, McLaughlin EA, Roman SD, Bromfield EG, Nixon B, et al. The impact of aging on macroautophagy in the pre-ovulatory mouse oocyte. Front Cell Dev Biol. 2021;9: 691826.

45. Yao Y, Wang B, Yu K, Song J, Wang L, Zhang X, et al. Nur77 improves ovarian function in reproductive aging mice by activating mitophagy and inhibiting apoptosis. Reprod Biol Endocrinol. 2024;22: 86.

46. Czarny P, Pawlowska E, Bialkowska-Warzecha J, Kaarniranta K, Blasiak J. Autophagy in DNA damage response. Int J Mol Sci. 2015;16: 2641–2662.

47. Gomes LR, Menck CFM, Leandro GS. Autophagy roles in the modulation of DNA repair pathways. Int J Mol Sci. 2017;18. doi:10.3390/ijms18112351

48. Nagy P, Sándor GO, Juhász G. Autophagy maintains stem cells and intestinal homeostasis in Drosophila. Sci Rep. 2018;8: 4644.

49. Simonsen A, Cumming RC, Brech A, Isakson P, Schubert DR, Finley KD. Promoting basal levels of autophagy in the nervous system enhances longevity and oxidant resistance in adult Drosophila. Autophagy. 2008;4: 176–184.

50. Damiano V, Spessotto P, Vanin G, Perin T, Maestro R, Santarosa M. The autophagy machinery contributes to E-cadherin turnover in breast cancer. Front Cell Dev Biol. 2020;8: 545.

